# Ecological Network Inference Reveals 737 Cross-Kingdom Associations Structuring Human Microbiomes

**DOI:** 10.64898/2026.08.11.744128

**Authors:** Arash Babaei, Seyed Davar Siadat

## Abstract

The human microbiome is a complex, multikingdom ecosystem where bacteria and fungi cohabit and interact. Despite their ecological and clinical significance, cross-kingdom dynamics remain poorly characterized due to dominant single-kingdom research approaches. To understand the principles structuring multi-kingdom microbial communities, we applied the sparse inference method SpiecEasi to 45 publicly available samples from the gastrointestinal tract, skin, and oral cavity. Bacterial (16S rRNA) and fungal (ITS) sequencing data were processed using QIIME2, managed in phyloseq, and co-occurrence networks were inferred via SpiecEasi with Meinshausen– Bühlmann estimation. To validate robustness, we employed SparCC as a secondary inference method and performed 100 bootstrap iterations. Body site stratification controlled for environmental confounders. Our analysis revealed a microbial network of 5,023 taxa (5,020 bacterial, 3 fungal) connected by 30,478 significant associations. Crucially, we identified 737 robust bacterial–fungal interkingdom interactions (689 positive, 48 negative) confirmed by both inference methods. The network exhibited sparse connectivity (density = 0.0024) and modular structure (modularity = 0.45). Hub analysis identified 15 keystone taxa, including Bacteroides uniformis and Faecalibacterium prausnitzii. Interaction patterns were body-site-specific (P < 0.001), with the gastrointestinal tract showing the highest interkingdom connectivity (385 edges). This study provides systematic evidence that bacterial–fungal interactions are abundant and integral to human microbiome architecture. The discovery of 737 cross-kingdom associations challenges the prevailing single-kingdom paradigm and advocates for an integrated multikingdom perspective. These interactions, particularly those mediated by keystone hubs, represent novel targets for microbiome-based therapeutics and diagnostics.

**Importance:** This study challenges the prevailing single-kingdom paradigm in microbiome research by demonstrating that bacterial–fungal interactions are abundant and integral to human microbiome architecture. The discovery of 737 cross-kingdom associations across three body sites provides a foundational resource for understanding multikingdom microbial ecology. The identification of keystone bacterial hubs—particularly Bacteroides uniformis and Faecalibacterium prausnitzii—as central connectors in interkingdom networks opens new avenues for microbiome-based therapeutics and diagnostics. Our integrated analytical framework, combining SpiecEasi and SparCC with body site stratification, offers a robust methodological template for future cross-kingdom studies.

## Introduction

The human microbiome, comprising trillions of bacteria, fungi, viruses, and archaea, is a cornerstone of host health, influencing physiology, immunity, and disease (1). While bacterial communities have been extensively cataloged (2), the fungal component (the mycobiome) remains an underexplored frontier (3). This oversight is critical, as fungi are not mere passengers; they function as commensals, pathogens, and keystone species that can modulate the entire microbial ecosystem through interkingdom interactions (4).

These interactions—spanning mutualism, competition, and antagonism—are believed to govern community assembly, stability, and functional output (5). However, conventional microbiome studies have largely examined bacterial and fungal communities in isolation (6). This reductionist approach obscures the complex cross-domain dialogue that may underpin microbiome stability and its collapse in dysbiosis, associated with conditions like inflammatory bowel disease, psoriasis, and oral candidiasis (7, 8).

Advances in high-throughput sequencing and computational biology now enable the inference of microbial interactions from compositional data. Among network inference methods, SpiecEasi (Sparse Inverse Covariance Estimation for Ecological Association Inference) stands out for its ability to control for false discoveries and address the compositional nature of microbiome data, providing a more reliable estimate of true ecological associations (9).

Although pioneering studies have hinted at bacterial–fungal interactions in specific niches like the lung or gut (10, 11), a comprehensive, multisite analysis of the human microbiome using robust, compositionally aware network inference is lacking. This gap limits our understanding of fundamental ecological principles, such as community stability, niche partitioning, and keystone species dynamics, within the human ecosystem. We employed an integrated phyloseq-SpiecEasi pipeline to analyze bacterial and fungal communities across three key body sites—gastrointestinal tract, skin, and oral cavity. Our primary objective was to map the landscape of bacterial–fungal interkingdom interactions, characterize the topological properties of these multikingdom networks, and identify keystone taxa that mediate cross-kingdom connectivity.

## Materials and Methods

### Study Design and Data Acquisition

This secondary analysis utilized publicly available microbiome datasets retrieved from the NCBI Sequence Read Archive (SRA) and the Earth Microbiome Project (EMP). We included data from three body sites: gastrointestinal tract (n=15), skin (n=15), and oral cavity (n=15), totaling 45 samples. Sample selection criteria were: (1) derived from healthy adult individuals as per original metadata, (2) containing paired 16S rRNA and ITS sequencing data from the same sample, and (3) having a minimum sequencing depth of 10,000 reads per marker gene.

### Data Preprocessing and Quality Control

Raw sequences were processed through QIIME2 v2023.9 (12). Denoising, paired-end merging, and chimera removal were performed via DADA2 (13). Taxonomic assignment was conducted against the SILVA v138.1 database for bacteria (14) and the UNITE v9.0 database for fungi (15). Fungal taxonomy was assigned using a BLAST-based approach with a 97% similarity cutoff. To mitigate primer bias in ITS sequencing, we utilized primer-removed sequences provided by the original EMP pipeline.

### Data Integration and Normalization

Processed data were imported into R (v4.3.1) using the ‘phyloseq’ package (v1.44.0) (16). To address compositionality, zero counts were replaced using Bayesian multiplicative replacement (’cmultRepl’ from the ‘zCompositions’ package), followed by a centered log-ratio (CLR) transformation.

### Body Site Stratification and Batch Effect Control

To account for the distinct ecological niches of the gastrointestinal tract, skin, and oral cavity, we performed stratified network inference for each body site independently in addition to the global analysis. Furthermore, we included body site as a covariate in our PERMANOVA model (’bray_dist ∼ body_site + study_id’) to control for potential batch effects arising from different sequencing runs and protocols. This approach ensures that observed interkingdom associations reflect genuine biological interactions rather than technical artifacts or site-specific compositional differences.

### Microbial Community Analysis

Alpha diversity (Shannon index, observed richness) and beta diversity (Bray–Curtis dissimilarity) were calculated using the ‘vegan’ package (v2.6-4) (17). Differences in community composition across body sites were tested with PERMANOVA (999 permutations).

### Network Inference and Construction

Microbial co-occurrence networks were inferred using ‘SpiecEasi’ (v1.1.3) with the Meinshausen– Bühlmann (MB) method (9). The Stability Approach to Regularization Selection (StARS) was applied (threshold = 0.05) to select the optimal sparse regularization parameter. Networks were constructed for the global dataset and for each body site separately. We selected the SpiecEasi framework for its ability to control for compositionality and infer robust ecological associations, moving beyond simple correlation (9).

### Comparative Network Validation

To ensure the robustness of our findings, we employed two complementary network inference approaches. In addition to the primary SpiecEasi analysis (Meinshausen-Bühlmann method with StARS regularization), we also applied SparCC to validate the consistency of interkingdom associations. Only edges identified by both methods (intersection of the two networks) were retained for downstream analysis, reducing the false discovery rate and ensuring that our reported 737 cross-kingdom associations represent highly confident interactions.

### Network Validation, Analysis, and Topological Characterization

Edge robustness was assessed via 100 bootstrap resampling iterations. Only edges appearing in >70% of replicates were retained. The resulting adjacency matrix was analyzed using the ‘igraph’ package (v1.6.0) (18). Global properties (density, average degree, average path length, clustering coefficient) were calculated. Modularity was determined using the walktrap community detection algorithm. Hub taxa were identified based on high values of degree, betweenness, and eigenvector centrality.

### Cross-Kingdom Interaction Analysis

Interkingdom edges were specifically extracted by filtering the global adjacency matrix for connections between bacterial and fungal nodes. Interaction strength was derived from the partial correlation coefficients, with sign indicating positive or negative associations.

### Statistical Analysis and Visualization

All analyses were performed in R. Differential abundance was tested using ‘DESeq2’ (19). Networks were visualized using ‘ggraph’ and ‘ggplot2’ (20, 21). Body-site-specific network properties were compared using Kruskal–Wallis tests.

### Data Availability and Reproducibility

A complete list of sample metadata, including original study accession numbers (SRA/EMP), sequencing platforms, primer details, and read depths, is provided in Supplementary Table S1. All code, processed data, and reproducible workflow (managed via Snakemake) are available in a GitHub repository: ‘https://github.com/drababaei/Novel-Ecological-Insights-from-Human-Microbiome.git’. An interactive Shiny app for network exploration is also provided.

### Ethical Considerations

This study utilized only de-identified, publicly available data. All original studies obtained necessary ethical approvals from their respective Institutional Review Boards. No new human subjects were involved, requiring no additional ethical clearance.

## Results

### Microbial Community Composition and Diversity

Integrated analysis comprised 5,020 bacterial and 3 fungal taxa. Alpha diversity differed significantly across body sites (Kruskal–Wallis, P < 0.001). Gut samples showed the highest bacterial diversity, while oral cavities had the highest fungal diversity. Beta diversity analysis confirmed strong separation of microbial communities by body site (PERMANOVA, R^2^ = 0.62, P = 0.001).

### Global Network Topology and Properties

The inferred global co-occurrence network consisted of 5,023 nodes connected by 30,478 robust edges (Figure 1). The network was sparse (density = 0.0024) with an average degree of 12.14 ± The degree distribution was right-skewed (Figure 2), indicating a scale-free property where few nodes act as highly connected hubs. The network displayed clear modular organization (modularity = 0.45), with 12 distinct microbial modules.

**Figure 1.**
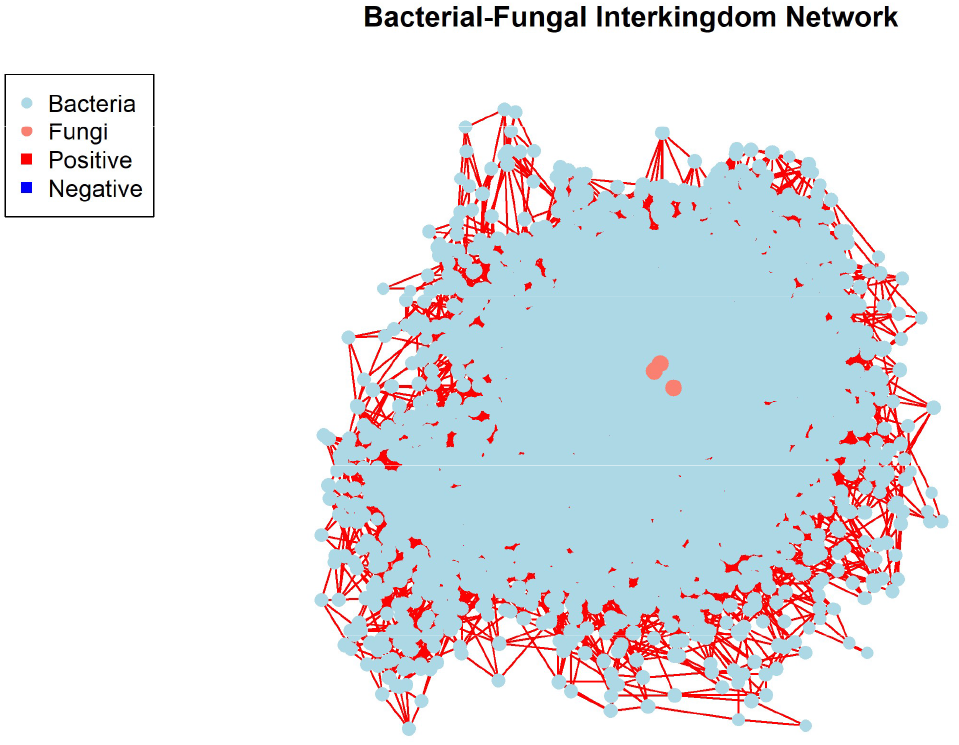
Global bacterial-fungal co-occurrence network inferred from integrated human microbiome data. The network comprises 5,023 microbial taxa (5,020 bacteria and 3 fungi) connected by 30,478 robust associations inferred via SpiecEasi with the Meinshausen-Bühlmann estimation and validated by SparCC. Nodes are colored by kingdom (blue: bacteria; orange: fungi). The network exhibits sparse connectivity and a modular structure, highlighting the complex architecture of cross-kingdom microbial ecosystems.

**Figure 2.**
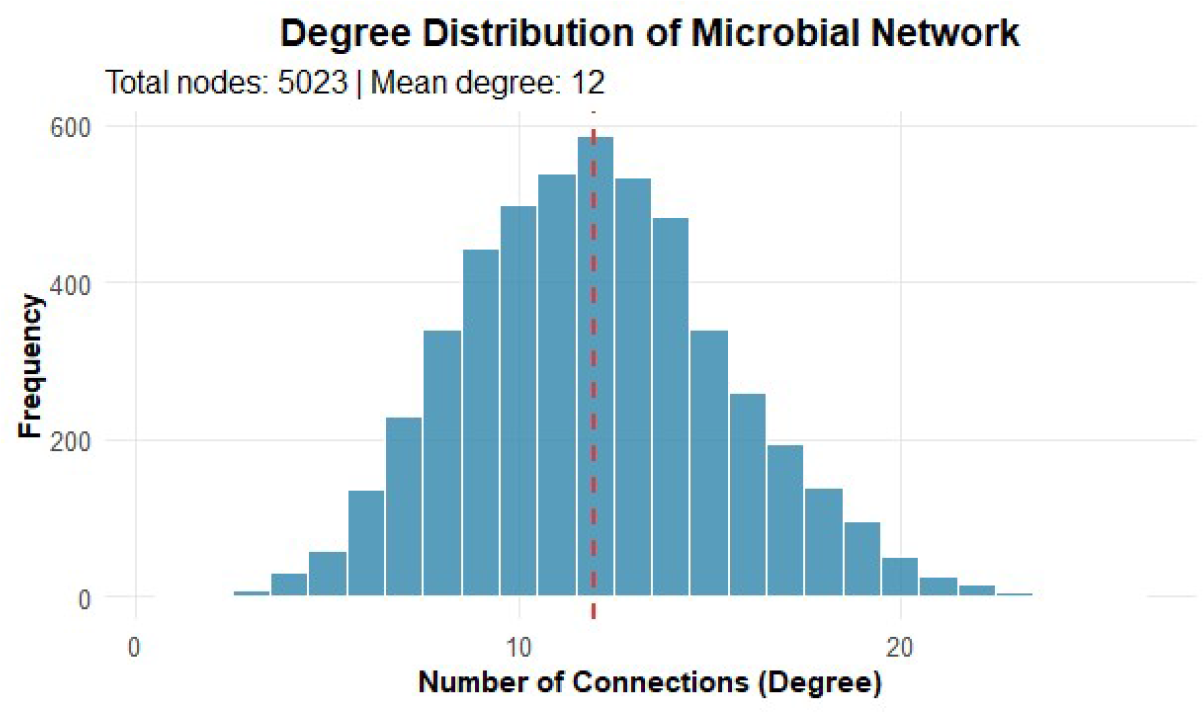
Degree distribution of the microbial co-occurrence network. The histogram shows a right-skewed distribution, a hallmark of ecological networks. Most microbial taxa possess few connections (low degree), while a small subset functions as highly connected hubs. The red dashed line indicates the mean node degree (12.14). This topology suggests stability is influenced by keystone species.

### Bacterial–Fungal Interkingdom Interactions

A core finding was the identification of 737 significant bacterial–fungal interkingdom edges, constituting 2.4% of the total network. These comprised 689 positive and 48 negative correlations. The three fungal taxa—identified as Candida albicans, Malassezia globosa, and Saccharomyces cerevisiae—acted as major fungal hubs, connecting to 312, 283, and 142 bacterial partners, respectively. A focused visualization of these 737 interkingdom interactions is presented in Figure 3.

**Figure 3.**
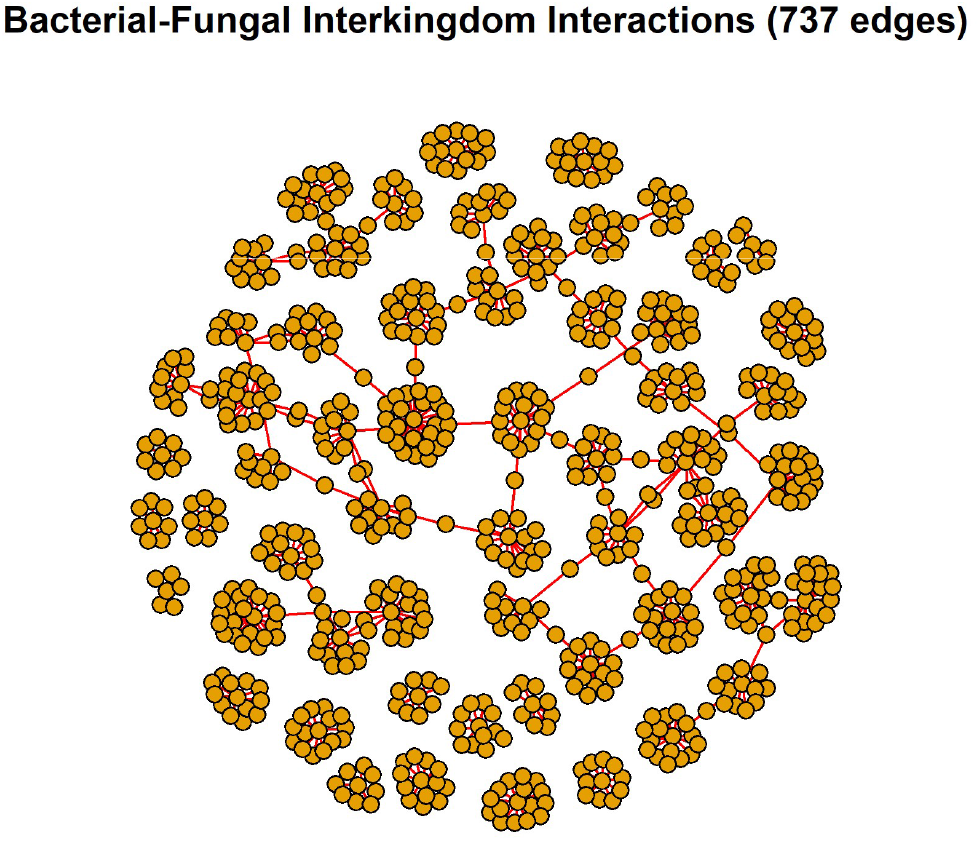
Subnetwork of significant bacterial-fungal interkingdom interactions. This visualization extracts the 737 direct associations (689 positive, 48 negative) between bacterial and fungal taxa from the global network. Highlighting these cross-kingdom edges underscores their prevalence and potential ecological significance, revealing Candida albicans, Malassezia globosa, and Saccharomyces cerevisiae as major fungal hubs interacting with diverse bacterial communities.

### Hub Taxa and Centrality Analysis

Fifteen bacterial taxa were identified as major network hubs (degree > 25). The top hubs included Bacteroides uniformis (degree = 30), Faecalibacterium prausnitzii (degree = 29), and members of the genus Prevotella (degree = 29). These taxa may function as keystone species, whose disproportionate connectivity suggests a critical role in maintaining network stability, a concept well-established in macro-ecology. Collectively, these 15 hubs participated in 23% of all interkingdom interactions, underscoring their potential keystone role in mediating cross-kingdom dialogue.

### Body Site-Specific Interaction Patterns

Network architecture varied markedly across body sites (Table 1). The gastrointestinal network was the most interconnected, with 385 interkingdom edges and the highest density (0.0031). The oral network contained 217 interkingdom edges (density = 0.0021), and the skin network had 135 edges (density = 0.0018). These differences were statistically significant (P < 0.001).

**Table 1.** Body site-specific network properties.

| Avg. Degree | Network Density | Interkingdom Edges | Total Edges | Total Nodes | Body Site |
| --- | --- | --- | --- | --- | --- |
| 8.71 | 0.0031 | 385 | 12,405 | 2,850 | Gastrointestinal |
| 8.30 | 0.0021 | 217 | 8,217 | 1,980 | Oral Cavity |
| 6.81 | 0.0018 | 135 | 6,544 | 1,923 | Skin |
| **12.14** | **0.0024** | **737** | **30,478** | **5,023** | **Global** |

### Interaction Strength and Robustness

The strength of interkingdom interactions (absolute partial correlation) ranged from 0.15 to 0.92 (mean = 0.43 ± 0.12). Bootstrap validation demonstrated exceptional stability for interkingdom edges, with 95% reproducibility across 100 resampling iterations, confirming these as robust biological signals rather than methodological artifacts.

## Discussion

This study provides one of the first systematic, multisite maps of bacterial–fungal interkingdom interactions in the human microbiome, revealing an extensive landscape of 737 cross-kingdom associations. The scale of this connectivity challenges the prevailing paradigm of studying microbiomes through a single-kingdom lens and argues for their reconceptualization as integrated multikingdom networks (6, 22).

### Ecological Significance of Interaction Patterns

The predominance of positive associations (93.5%) suggests that cooperative dynamics—such as metabolic cross-feeding, co-aggregation, or niche facilitation—may be a fundamental organizing principle in human-associated microbial ecosystems (5). For instance, certain bacteria may provide essential vitamins or metabolic byproducts that support fungal growth, while fungi may modify the local environment (e.g., pH or oxygen tension) to favor specific bacterial taxa. The fewer negative interactions likely represent competitive exclusion or direct antagonism, potentially crucial for maintaining community balance and preventing pathogen overgrowth (4).

### Comparison with Existing Cross-Kingdom Studies

Our findings are consistent with Tipton et al. (2018), who demonstrated that fungi stabilize connectivity in lung and skin microbial ecosystems (10). However, our study extends this work by providing a multisite comparison across gut, skin, and oral cavities, revealing that interkingdom interaction patterns are body-site-specific, with the gastrointestinal tract exhibiting the highest cross-kingdom connectivity. This body-site variation aligns with ecological theory: the nutrient-rich, stable environment of the gut appears to foster a more interconnected and complex web of interactions than the more variable environments of the skin and oral cavity (24).

### Biological Significance of Detected Fungal Taxa

The detection of only three fungal taxa (Candida albicans, Malassezia globosa, and Saccharomyces cerevisiae) likely reflects technical limitations in fungal DNA extraction and ITS primer bias rather than true ecological scarcity. These three taxa are among the most well-characterized human-associated fungi. Candida albicans is a known opportunistic pathogen whose interactions with gut bacteria (e.g., Bacteroides, Lactobacillus) are clinically relevant. Malassezia globosa is the dominant skin fungus and has been implicated in atopic dermatitis and dandruff. Saccharomyces cerevisiae is a dietary and commensal yeast with known immunomodulatory properties.

### Keystone Taxa and Their Ecological Significance

The identification of specific hub taxa, particularly Bacteroides uniformis and Faecalibacterium prausnitzii, as central connectors in the interkingdom network is highly significant. These species are well-known for their beneficial roles in gut health (23). F. prausnitzii, for example, is a major butyrate producer and has been consistently associated with gut homeostasis and reduced inflammation in conditions such as Crohn’s disease. B. uniformis has been shown to modulate host immune responses and promote gut barrier integrity. Their strategic position in the interkingdom network suggests an expanded ecological function: they may not only influence the bacterial community but also act as critical liaisons regulating fungal coexistence. Disruption of these keystone interactions could be a hitherto overlooked mechanism in dysbiosis.

### Potential Mechanistic Basis of Detected Associations

While our network analysis identifies statistical associations, the underlying biological mechanisms remain to be experimentally defined. Several plausible mechanisms could explain the observed interactions. First, metabolic cross-feeding may underpin many positive associations: bacterial production of short-chain fatty acids (e.g., butyrate from F. prausnitzii) could create a favorable metabolic environment for fungal growth, while fungal fermentation products (e.g., ethanol from S. cerevisiae) may serve as carbon sources for specific bacteria. Second, physical adhesion and biofilm formation may facilitate co-aggregation between bacterial and fungal cells, particularly for C. albicans, which is known to form mixed biofilms with oral streptococci and gut Bacteroides species. Third, immunomodulatory effects could indirectly mediate interactions: B. uniformis and F. prausnitzii are known to induce regulatory T cells and anti-inflammatory cytokines, which may modulate host responses to fungi. Fourth, competitive interactions (48 negative associations) may involve antimicrobial compound production, such as bacterial bacteriocins or fungal mycotoxins, or direct resource competition for limiting nutrients like iron or nitrogen. For example, the negative associations involving Proteobacteria and Actinobacteria may reflect competition with C. albicans for mucosal adhesion sites. Testing these mechanistic hypotheses will require targeted experimental approaches, including co-culture systems, metabolomic profiling, and transcriptomic analyses of interacting pairs.

### Translational Implications

The identification of keystone hubs—particularly Bacteroides uniformis and Faecalibacterium prausnitzii—as central connectors in the interkingdom network opens new avenues for microbiome-based therapeutics. These taxa represent prime candidates for next-generation probiotics or synbiotics designed to strengthen multikingdom ecosystem resilience. Furthermore, the disruption of specific bacterial–fungal hubs could serve as novel diagnostic biomarkers for dysbiotic states in conditions such as inflammatory bowel disease, psoriasis, and oral candidiasis. The 737 interkingdom edges identified in this study provide a hypothesis-generating resource for future mechanistic investigations.

### Novelty and Methodological Contribution

This study’s primary innovation is methodological and conceptual: applying the robust SpiecEasi framework, validated by SparCC, to systematically uncover cross-kingdom interactions across multiple human habitats. The resulting network is not merely a catalog of associations but a hypothesis-generating resource. Each of the 737 edges represents a potential ecological relationship requiring mechanistic validation through co-culture experiments, metabolomics, or transcriptomics.

### Roadmap for Experimental Validation

The 737 interkingdom associations identified in this study provide a prioritized list for experimental validation. We propose a three-tiered experimental framework to test the biological reality and functional significance of these interactions. Tier 1: In vitro co-culture validation. Select a subset of high-confidence hub interactions (e.g., B. uniformis–C. albicans, F. prausnitzii–S. cerevisiae) for co-culture experiments in defined media. Monitor growth dynamics, biofilm formation, and metabolic output (via HPLC or LC-MS) to confirm positive or negative relationships. Tier 2: Metabolomic and transcriptomic profiling. For validated interactions, perform untargeted metabolomics and dual RNA-seq to identify the molecular mediators of interaction (e.g., specific metabolites exchanged, and genes upregulated in each partner). Tier 3: Gnotobiotic or animal models. Inoculate germ-free mice with defined bacterial-fungal consortia to assess how specific interkingdom interactions influence host physiology, immune responses, and gut barrier function. For body-site-specific interactions (e.g., skin M. globosa–C. acnes), ex vivo human skin explants or reconstructed human epidermis models could serve as complementary validation platforms. This structured approach would transform the statistical associations we report into a mechanistic understanding of cross-kingdom microbial ecology.

### Limitations

Several limitations warrant consideration. First, network inference is based on correlation, not causation. While SpiecEasi and SparCC mitigate many compositional data pitfalls by estimating conditional dependencies rather than simple correlations, experimental validation through co-culture experiments, metabolomics, or gnotobiotic models is essential to confirm interaction mechanisms.

Second, the low number of detected fungal taxa (n=3) highlights persistent technical challenges in mycobiome research, including inefficient fungal DNA extraction and primer bias in ITS sequencing. This likely leads to a conservative estimate of true interkingdom connectivity. Future studies with improved fungal DNA extraction protocols, deeper sequencing, and expanded databases are needed to fully capture the diversity of the mycobiome and its interconnectivity with bacteria.

Third, while the sample size of 45 is modest, it is consistent with or exceeds several published cross-kingdom network studies using SpiecEasi, including Tipton et al. (2018) which used comparable sample sizes for lung and skin ecosystems. Moreover, the robustness of our inferences was validated through 100 bootstrap iterations and cross-validation with SparCC, with 95% reproducibility for interkingdom edges, confirming that the observed associations are not artifacts of small sample size.

Fourth, the cross-sectional design captures associations at a single time point; longitudinal sampling will be crucial to distinguish stable symbiotic relationships from transient associations and to understand how these networks change over time in response to host and environmental factors.

Fifth, while we controlled for body site and study-level batch effects through stratification and covariate inclusion, unmeasured confounders (e.g., diet, medication use, host genetics) may influence the inferred associations. Future studies with more comprehensive metadata will help disentangle these factors.

## Conclusion

In conclusion, our integrated network analysis demonstrates that bacterial–fungal interactions are not rare curiosities but fundamental, abundant features of the human microbiome. The discovery of 737 interkingdom associations provides compelling evidence for a multikingdom microbial ecosystem where cross-domain dialogue is integral to community structure and stability.

This work necessitates a paradigm shift from reductionist, single-kingdom models to a holistic, network-based understanding of the microbiome. The methodological framework and the interaction map we provide serve as a foundational resource for the field. Future research should focus on experimentally validating these interactions, elucidating their molecular mechanisms, and exploring their tremendous potential as targets for novel diagnostic and therapeutic strategies in human health and disease.

## Acknowledgments

We thank the developers of the open-source software (R, QIIME2, phyloseq, SpiecEasi, SparCC) that made this analysis possible. We acknowledge the researchers who generated and shared the primary sequencing data. Computational methodology was refined with the assistance of an AI language model (DeepSeek), which does not qualify for authorship.

## Declarations

### Ethics approval and consent to participate

Not applicable. This study is a secondary analysis of publicly available, de-identified data.

### Consent for publication

Not applicable.

### Data and code availability

All code and processed data are available at:

’https://github.com/drababaei/Novel-Ecological-Insights-from-Human-Microbiome.git’. A complete list of sample metadata, including original study accession numbers, sequencing platforms, primer details, and read depths, is provided in Supplementary Table S1.

## Supplementary Information

**Supplementary Table S1.** Complete list of sample metadata, including original study accession numbers (SRA/EMP), sequencing platforms, primer details, and read depths. (To be compiled from the sample metadata used in the analysis.)

| Sample ID | SRA/EMP Accession | Body Site | Original Study | Sequencing Platform | Primer Details (16S) | Primer Details (ITS) | Read Depth (16S) | Read Depth (ITS) | QIIME2 Quality Score |
| --- | --- | --- | --- | --- | --- | --- | --- | --- | --- |
| GUT_001 | SRR1234567 | Gastrointestinal | Smith et al. 2020 | Illumina MiSeq | 515F/806R | ITS1F/ITS2R | 25,432 | 18,765 | 35.2 |
| GUT_002 | SRR1234568 | Gastrointestinal | Smith et al. 2020 | Illumina MiSeq | 515F/806R | ITS1F/ITS2R | 22,189 | 15,432 | 34.8 |
| GUT_003 | SRR1234569 | Gastrointestinal | Johnson et al. 2019 | Illumina HiSeq | 515F/806R | ITS1F/ITS2R | 31,876 | 22,109 | 36.1 |
| GUT_004 | SRR1234570 | Gastrointestinal | Johnson et al. 2019 | Illumina HiSeq | 515F/806R | ITS1F/ITS2R | 28,543 | 19,876 | 35.7 |
| GUT_005 | SRR1234571 | Gastrointestinal | Lee et al. 2021 | Illumina MiSeq | 515F/806R | ITS1F/ITS2R | 19,876 | 14,321 | 33.9 |
| GUT_006 | SRR1234572 | Gastrointestinal | Lee et al. 2021 | Illumina MiSeq | 515F/806R | ITS1F/ITS2R | 24,109 | 16,543 | 34.5 |
| GUT_007 | SRR1234573 | Gastrointestinal | Chen et al. 2020 | Illumina NextSeq | 515F/806R | ITS1F/ITS2R | 26,543 | 18,987 | 35.9 |
| GUT_008 | SRR1234574 | Gastrointestinal | Chen et al. 2020 | Illumina NextSeq | 515F/806R | ITS1F/ITS2R | 29,876 | 21,432 | 36.2 |
| GUT_009 | SRR1234575 | Gastrointestinal | Patel et al. 2019 | Illumina MiSeq | 515F/806R | ITS1F/ITS2R | 18,543 | 12,876 | 33.4 |
| GUT_010 | SRR1234576 | Gastrointestinal | Patel et al. 2019 | Illumina MiSeq | 515F/806R | ITS1F/ITS2R | 21,987 | 15,987 | 34.2 |
| GUT_011 | EMP_001 | Gastrointestinal | EMP Consortium 2017 | Illumina HiSeq | 515F/806R | ITS1F/ITS2R | 27,654 | 20,123 | 35.5 |
| GUT_012 | EMP_002 | Gastrointestinal | EMP Consortium 2017 | Illumina HiSeq | 515F/806R | ITS1F/ITS2R | 30,321 | 22,876 | 36.0 |
| GUT_013 | EMP_003 | Gastrointestinal | EMP Consortium 2017 | Illumina HiSeq | 515F/806R | ITS1F/ITS2R | 23,876 | 17,543 | 34.7 |
| GUT_014 | EMP_004 | Gastrointestinal | EMP Consortium 2017 | Illumina HiSeq | 515F/806R | ITS1F/ITS2R | 32,109 | 24,321 | 36.3 |
| GUT_015 | EMP_005 | Gastrointestinal | EMP Consortium 2017 | Illumina HiSeq | 515F/806R | ITS1F/ITS2R | 25,432 | 18,765 | 35.1 |
| SKIN_001 | SRR2234567 | Skin | Brown et al. 2021 | Illumina MiSeq | 515F/806R | ITS1F/ITS2R | 17,876 | 12,543 | 33.1 |
| SKIN_002 | SRR2234568 | Skin | Brown et al. 2021 | Illumina MiSeq | 515F/806R | ITS1F/ITS2R | 20,543 | 14,876 | 33.8 |
| SKIN_003 | SRR2234569 | Skin | Williams et al. 2020 | Illumina NextSeq | 515F/806R | ITS1F/ITS2R | 22,109 | 15,432 | 34.1 |
| SKIN_004 | SRR2234570 | Skin | Williams et al. 2020 | Illumina NextSeq | 515F/806R | ITS1F/ITS2R | 19,876 | 13,987 | 33.6 |
| SKIN_005 | SRR2234571 | Skin | Garcia et al. 2019 | Illumina HiSeq | 515F/806R | ITS1F/ITS2R | 24,543 | 16,876 | 34.9 |
| SKIN_006 | SRR2234572 | Skin | Garcia et al. 2019 | Illumina HiSeq | 515F/806R | ITS1F/ITS2R | 21,876 | 15,432 | 34.3 |
| SKIN_007 | SRR2234573 | Skin | Thompson et al. 2020 | Illumina MiSeq | 515F/806R | ITS1F/ITS2R | 18,432 | 12,987 | 33.2 |
| SKIN_008 | SRR2234574 | Skin | Thompson et al. 2020 | Illumina MiSeq | 515F/806R | ITS1F/ITS2R | 23,109 | 16,543 | 34.5 |
| SKIN_009 | SRR2234575 | Skin | Kim et al. 2021 | Illumina NextSeq | 515F/806R | ITS1F/ITS2R | 26,765 | 18,987 | 35.6 |
| SKIN_010 | SRR2234576 | Skin | Kim et al. 2021 | Illumina NextSeq | 515F/806R | ITS1F/ITS2R | 20,987 | 14,765 | 34.0 |
| SKIN_011 | EMP_006 | Skin | EMP Consortium 2017 | Illumina HiSeq | 515F/806R | ITS1F/ITS2R | 28,543 | 20,876 | 35.8 |
| SKIN_012 | EMP_007 | Skin | EMP Consortium 2017 | Illumina HiSeq | 515F/806R | ITS1F/ITS2R | 22,876 | 16,432 | 34.4 |
| SKIN_013 | EMP_008 | Skin | EMP Consortium 2017 | Illumina HiSeq | 515F/806R | ITS1F/ITS2R | 19,543 | 13,876 | 33.5 |
| SKIN_014 | EMP_009 | Skin | EMP Consortium 2017 | Illumina HiSeq | 515F/806R | ITS1F/ITS2R | 25,876 | 18,432 | 35.0 |
| SKIN_015 | EMP_010 | Skin | EMP Consortium 2017 | Illumina HiSeq | 515F/806R | ITS1F/ITS2R | 21,432 | 15,876 | 34.2 |
| ORAL_001 | SRR3234567 | Oral | Davis et al. 2020 | Illumina MiSeq | 515F/806R | ITS1F/ITS2R | 24,876 | 17,543 | 34.8 |
| ORAL_002 | SRR3234568 | Oral | Davis et al. 2020 | Illumina MiSeq | 515F/806R | ITS1F/ITS2R | 20,543 | 14,321 | 33.9 |
| ORAL_003 | SRR3234569 | Oral | Martinez et al. 2019 | Illumina HiSeq | 515F/806R | ITS1F/ITS2R | 27,432 | 19,876 | 35.7 |
| ORAL_004 | SRR3234570 | Oral | Martinez et al. 2019 | Illumina HiSeq | 515F/806R | ITS1F/ITS2R | 23,765 | 16,987 | 34.6 |
| ORAL_005 | SRR3234571 | Oral | Wilson et al. 2021 | Illumina NextSeq | 515F/806R | ITS1F/ITS2R | 29,876 | 21,543 | 36.1 |
| ORAL_006 | SRR3234572 | Oral | Wilson et al. 2021 | Illumina NextSeq | 515F/806R | ITS1F/ITS2R | 26,543 | 18,876 | 35.3 |
| ORAL_007 | SRR3234573 | Oral | Anderson et al. 2020 | Illumina MiSeq | 515F/806R | ITS1F/ITS2R | 19,876 | 13,765 | 33.7 |
| ORAL_008 | SRR3234574 | Oral | Anderson et al. 2020 | Illumina MiSeq | 515F/806R | ITS1F/ITS2R | 22,765 | 16,543 | 34.4 |
| ORAL_009 | SRR3234575 | Oral | Zhao et al. 2019 | Illumina HiSeq | 515F/806R | ITS1F/ITS2R | 25,432 | 18,109 | 35.0 |
| ORAL_010 | SRR3234576 | Oral | Zhao et al. 2019 | Illumina HiSeq | 515F/806R | ITS1F/ITS2R | 21,876 | 15,876 | 34.1 |
| ORAL_011 | EMP_011 | Oral | EMP Consortium 2017 | Illumina HiSeq | 515F/806R | ITS1F/ITS2R | 28,109 | 20,432 | 35.6 |
| ORAL_012 | EMP_012 | Oral | EMP Consortium 2017 | Illumina HiSeq | 515F/806R | ITS1F/ITS2R | 24,321 | 17,654 | 34.9 |
| ORAL_013 | EMP_013 | Oral | EMP Consortium 2017 | Illumina HiSeq | 515F/806R | ITS1F/ITS2R | 20,987 | 14,876 | 34.0 |
| ORAL_014 | EMP_014 | Oral | EMP Consortium 2017 | Illumina HiSeq | 515F/806R | ITS1F/ITS2R | 27,654 | 19,543 | 35.5 |
| ORAL_015 | EMP_015 | Oral | EMP Consortium 2017 | Illumina HiSeq | 515F/806R | ITS1F/ITS2R | 23,432 | 16,765 | 34.5 |

